# Protein kinase G inhibition preserves photoreceptor viability and function in a new mouse model for autosomal dominant retinitis pigmentosa

**DOI:** 10.1101/2024.12.20.629748

**Authors:** Yu Zhu, Lucia Peiroten, Pranav Nanda Kumar, Catherine Hottin, Wadood Haq, François Paquet-Durand

## Abstract

Retinitis Pigmentosa (RP) is the most common inherited retinal degeneration, characterized by an initial loss of rod photoreceptor cells. Photoreceptor cell death has been associated with high levels of cyclic guanosine-3′, 5′-monophosphate (cGMP) in animal models of autosomal recessive RP (ARRP) and autosomal dominant RP (ADRP). cGMP analogues inhibiting protein kinase G (PKG) have been found to prevent rod degeneration in ARRP disease models, but their effects on ADRP are unknown. Here, we used the recently generated rhodopsin-mutant *Rho*^I255d/+^ ADRP mouse model to investigate cGMP-signaling and the effects of cGMP analogues targeting PKG.

cGMP accumulation was investigated by retinal immunostaining in wild-type (WT), *Rho*^I255d/+^, and *Rho*^I255d/I255d^ mice. The therapeutic efficacy of the cGMP analogues CN03 and CN238 was evaluated on organotypic retinal explant cultures derived from WT and *Rho*^I255d/+^ mice. Readouts included the TUNEL assay and immunostaining for cone arrestin-3. Downstream effectors of cell death were visualized using calpain, poly-ADP-ribose polymerase (PARP), and histone deacetylase (HDAC) *in situ* assays, as well as caspase-3 immunostaining. Photoreceptor function was assessed using micro-electroretinogram (µERG) recordings. When compared with WT, *Rho*^I255d^ photoreceptors displayed cGMP accumulation in outer segments. In the *Rho*^I255d/+^ ADRP model, CN03 and CN238 significantly reduced the number of dying photoreceptors. However, the relatively small number of photoreceptors exhibiting caspase-3 activity was not changed by the treatment. Remarkably, CN238 effectively provided long-lasting neuroprotection of cone photoreceptors and preserved retinal light responsiveness of *Rho*^I255d/+^ retina.

Overall, this study suggests caspase-independent but cGMP-dependent cell death as a dominant degenerative mechanism in the *Rho*^I255d/+^ ADRP mouse model. PKG inhibition demonstrated robust neuroprotection of both rod and cone photoreceptors, while the marked preservation of retinal function, especially with the compound CN238, highlighted cGMP analogues for the treatment of ADRP.

## Introduction

Retinitis pigmentosa (RP) is a blinding disease characterized by the primary cell death of rod photoreceptors, followed by secondary loss of cone photoreceptors. RP affects approximately 1 in 4 000 people worldwide (1). Mutations in over 80 RP disease genes can be categorized by three main inheritance patterns: Autosomal recessive (ARRP; 50-60% of all cases), autosomal dominant (ADRP; 30-40%), and X-linked (5-15%) (2). Disease-causing variants of the rhodopsin gene (*RHO*) are most commonly associated with ADRP, with more than 150 documented missense or nonsense *RHO* mutations (3).

The mechanisms underlying cell death in RP are still largely unclear. However, high levels of cyclic guanosine-3′, 5′-monophosphate (cGMP) are known to play a critical role in various ARRP animal models, such as in *rd1* and *rd10* mice, as well as in *Rho*^P23H^ and *Rho*^S334ter^ ADRP rat models (4). Elevated cGMP levels are often caused by mutations that disrupt the phototransduction cascade, and may drive photoreceptor degeneration through two distinct pathways: 1) Activation of protein kinase G (PKG), likely connected to histone deacetylase (HDAC) and poly-ADP-ribose polymerase (PARP) activity (4, 5). 2) Activation of cyclic nucleotide-gated (CNG) -channels, with the resultant Ca^2+^ influx associated with the activation of Ca^2+^-dependent, calpain-type proteases (6). However, CNG-channels activity is essential for visual phototransduction and compounds inhibiting CNG-channels likely are detrimental to photoreceptor viability (7). Thus, to develop new drugs for RP treatment, analogues of cGMP have been designed to specifically inhibit PKG. These compounds bear a sulfur atom to form Rp-configurated phosphorothioates (Rp-cGMPS), which bind to PKG without activating the kinase (8, 9). Inhibition of PKG previously demonstrated photoreceptor neuroprotection in *rd1*, *rd2*, and *rd10* ARRP mouse models (10, 11).

A case of *RHO*-associated ADRP was originally identified in a British family carrying an in-frame 3-bp deletion at codons 255/256 of *RHO*. This results in the loss of one isoleucine (I) residue in rhodopsin protein and causes severe rod degeneration early in life (12). The human homologous *Rho*^I255d^ knock-in mouse model was recently generated to allow for pertinent ADRP research (13). Notably, in *Rho*^I255d^ retina, a rapid loss of photoreceptors was associated with increased activity of calpain and PARP (13), suggesting that the underlying cell death mechanism was linked to cGMP-signaling (5).

To study the possible involvement of cGMP-dependent cell death in ADRP caused by *RHO* mutations, we investigated cGMP accumulation in *Rho*^I255d^ photoreceptors and evaluated the neuroprotective effect of cGMP analogues on organotypic retinal explant cultures derived from heterozygous *Rho*^I255d/+^ mice. Remarkably, the novel PKG inhibitor CN238 – a cGMP analogue – demonstrated long-lasting neuroprotection of both rod and cone photoreceptors, preserving retinal function in the long-term. These findings reveal excessive cGMP-signaling as a major disease driver in ADRP and highlight the potential of cGMP analogues for ADRP therapy development.

## Results

### cGMP accumulates in the outer segments of *Rho*^l255d/+^ and *Rho*^l255d/I255d^ photoreceptors

High intracellular cGMP levels have been observed in the photoreceptors of various RP animal models, including ADRP rat models (4, 14). Previous work on the *Rho*^I255d^ mutation in heterozygous and homozygous mice found that photoreceptor cell death peaks at or just before postnatal day (P) 20 (13). Hence, to assess the possible role of cGMP in *Rho*^I255d^ mutation-dependent cell death, we performed immunostaining on P20 retinal cross-sections obtained from heterozygous *Rho*^I255d/+^ (*R*^I255d/+^), homozygous *Rho*^I255d/I255d^ (*R*^I255d/I255d^), or wild-type (WT) mice (Fig. 1).

**Figure 1.**
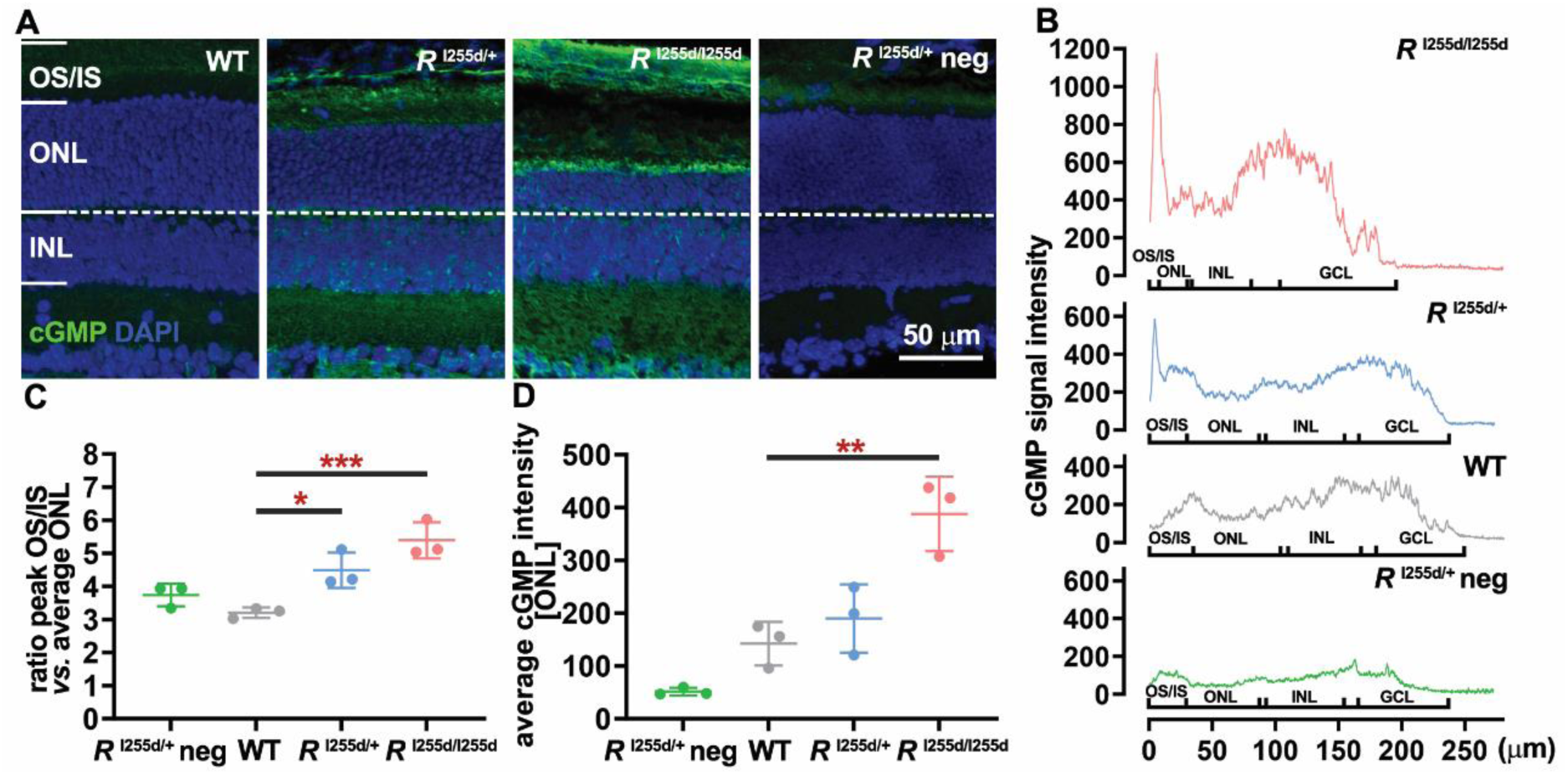
cGMP accumulation in degenerating *Rho*^I255d^ mutant photoreceptors. **A.** At post-natal day (P) 20, cGMP immunostaining (green) labeled photoreceptor outer and inner segments (OS/IS) of heterozygous *Rho*^I255d/+^ and homozygous *Rho*^I255d/I255d^ retina. Such a label was not seen in the wild-type (WT) and *Rho*^I255d/+^ negative control (neg.; without cGMP-antibody). DAPI (blue) was used as nuclear counterstain. **B.** Quantification of cGMP-staining intensity across retinal layers. Marked differences in cGMP staining intensity were seen at the level of OS/IS. **C.** Ratios of cGMP-staining intensity, comparing OS/IS with outer nuclear layer (ONL). **D.** Average cGMP staining intensity in ONL. Images represent results obtained from three different animals; error bars indicate SD; statistical testing: One-way ANOVA followed by Dunnett’s multiple comparisons test; significance levels: * p ≤ 0.05, ** p ≤ 0.01, *** p ≤ 0.001, INL = inner nuclear layer; scale bar = 50 µm.

cGMP labeling was detected in the outer and inner segments (OS/IS) in *Rho*^I255d/+^ and *Rho*^I255d/I255d^ mice, whereas no obvious signal was found in WT or in the negative control retina (Fig. 1A). The overall cGMP signal intensity across the retina was elevated in the mutant OS/IS (Fig. 1B, Table 1). The ratio of cGMP peak staining intensity in OS/IS vs. average staining intensity in the outer nuclear layer (ONL) was significantly higher in both *Rho*^I255d/+^ and *Rho*^I255d/I255d^ mice, compared with WT (Fig. 1C, Table 2). Additionally, the average cGMP intensity in the ONL was significantly increased in *Rho*^I255d/I255d^ mice compared with WT (Fig. 1D, Table 2). These findings indicated cGMP accumulation in the *Rho*^I255d^ retina, suggesting a possible causative role in retinal degeneration in this new ADRP mouse model.

### Elevated calpain-2 activation is linked to retinal degeneration in *Rho*^l255d/+^ retina

To represent the human clinical genotype, further investigations focused on the heterozygous *Rho*^I255d/+^ mutant (12). *Rho*^I255d/+^ retina displays progressive photoreceptor loss, which at P20 was linked to increased calpain activity in the ONL (13). Most of this calpain activity appears to come from calpain-2, a calpain isoform activated by millimolar Ca^2+^, that has been implicated repeatedly in retinal degeneration (15, 16). To link calpain-2 with *Rho*^I255d/+^ retinal degeneration, we performed TUNEL assays to label dying cells and immunostaining for activated calpain-2 on retinal sections derived from *Rho*^I255d/+^ and WT at P20 (Fig. 2). The percentage of TUNEL-positive cells in the ONL of *Rho*^I255d/+^ retina was significantly increased when compared to WT (Fig. 2A, B, Table 3). Similarly, immunolabelling for calpain-2 revealed significantly increased numbers of ONL cells (Fig. 2C, D, Table 3). Still, the absolute numbers of calpain-2 positive cells were lower compared to TUNEL positive cell numbers. These results indicated that calpain-2 activity was connected with *Rho*^I255d/+^ photoreceptor cell death.

**Figure 2.**
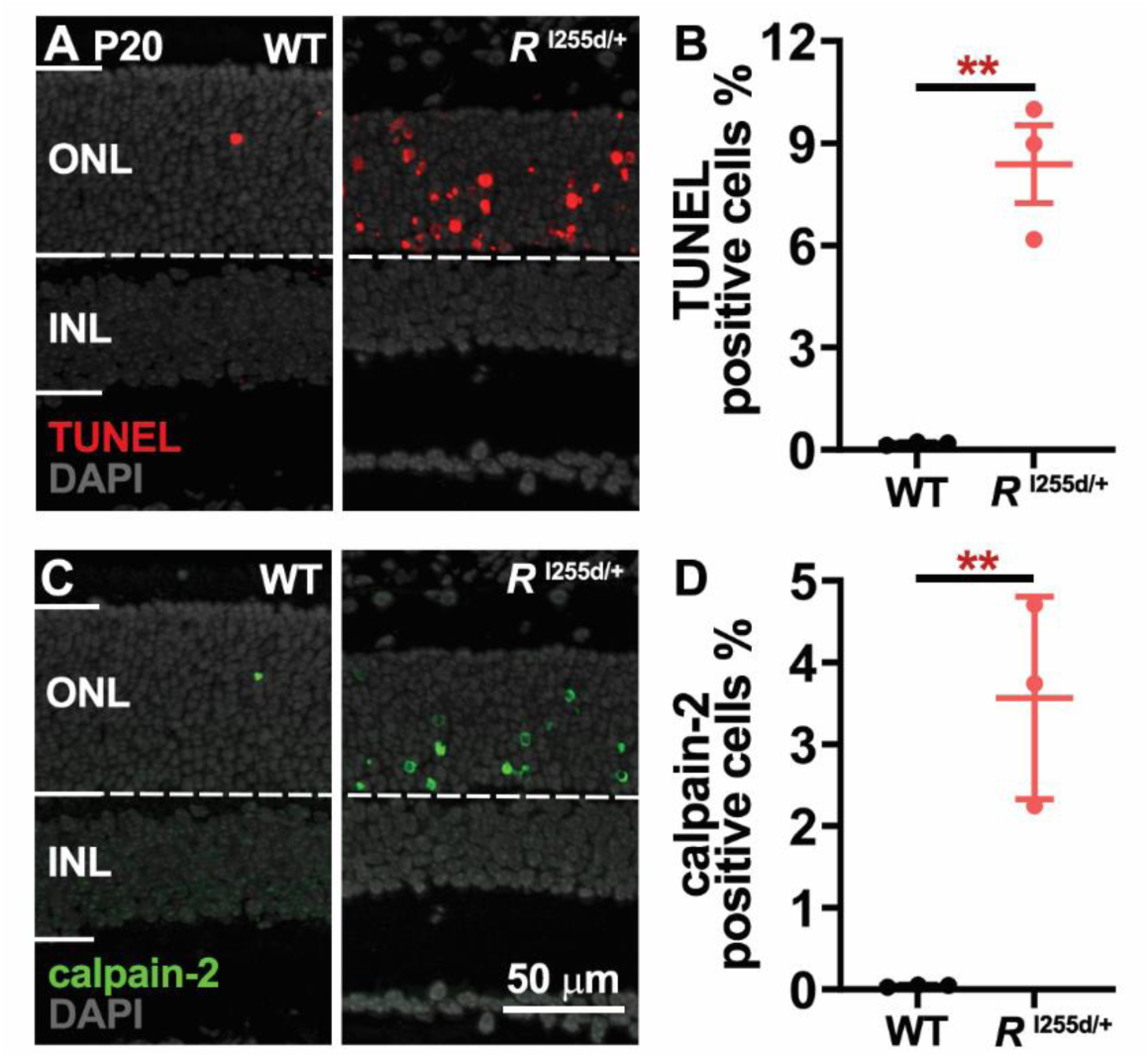
Increased calpain-2 activation correlates with *Rho*^I255d/+^ photoreceptor death. **A.** Photoreceptor cell death in the outer nuclear layer (ONL) was visualized via TUNEL assay (red) in wild-type (WT) and *Rho*^I255d/+^ retina at post-natal (P) day 20. **B.** Quantification of TUNEL-positive cells in ONL showed a significant increase in *Rho*^I255d/+^ compared with WT. **C.** Immunostaining of calpain-2 (green) in WT and *Rho*^I255d/+^ revealed increased activation in the mutant. **D.** The percentage of calpain-2-positive cells in ONL confirmed higher calpain-2 activation in *Rho*^I255d/+^ compared with WT. DAPI (grey) was used as nuclear counterstain. Images represent results obtained from three animals; error bars indicate SD; statistical testing: unpaired *t*-test; ** p ≤ 0.01; INL = Inner nuclear layer; scalar bar = 50 µm.

### cGMP analogues preserve *Rho*^I255d/+^ photoreceptors in short- and long-term treatments

To explore the possible involvement of cGMP-dependent PKG in *Rho*^I255d/+^ retinal degeneration, we employed the selective inhibitory cGMP analogues CN03 and CN238 (11). Organotypic retinal explants derived from *Rho*^I255d/+^ and WT mice were cultured and maintained under entirely controlled conditions – free of serum and antibiotics – from P12 to P18 and P20 (short-term), or P24 and P28 (long-term) and exposed to 50 μM of either cGMP analogue. To allow retinas to adapt to culturing conditions treatments started two days after explantation, i.e. at P14. To assess the effect of cGMP analogue on retinal degeneration in *Rho*^I255d/+^ retina, cell death was quantified via TUNEL assays in treated and non-treated (NT) retinas.

Increased numbers of TUNEL-positive cells were observed NT mutant retinas compared with WT (Fig. 3). Remarkably, the PKG inhibitors CN03 and CN238 significantly decreased photoreceptor cell death in short- and long-term retinal explant cultures compared with NT mutant (Fig. 3B, E, H, K, Table 4). Between P18 and P28 the numbers of surviving photoreceptors, quantified as ONL cell rows, significantly decreased in NT mutant retina when compared to WT (Fig. 3C, F, I, L). In short-term cultures ending at P18 and P20, *i.e.* at time-points at which the *Rho*^I255d/+^ degeneration has not yet produced a major loss of photoreceptor cells, the ONL row counts in CN03 or CN238 treated mutant retina did not significantly differ from NT (Fig. 3C, F). However, in long-term cultures lasting until P24 and P28 CN238 treatment resulted in a significant preservation of ONL rows in *Rho*^I255d/+^ retina. These data suggest that cGMP analogues, particularly CN238, can effectively reduce *Rho*^I255d/+^ photoreceptor degeneration.

**Figure 3.**
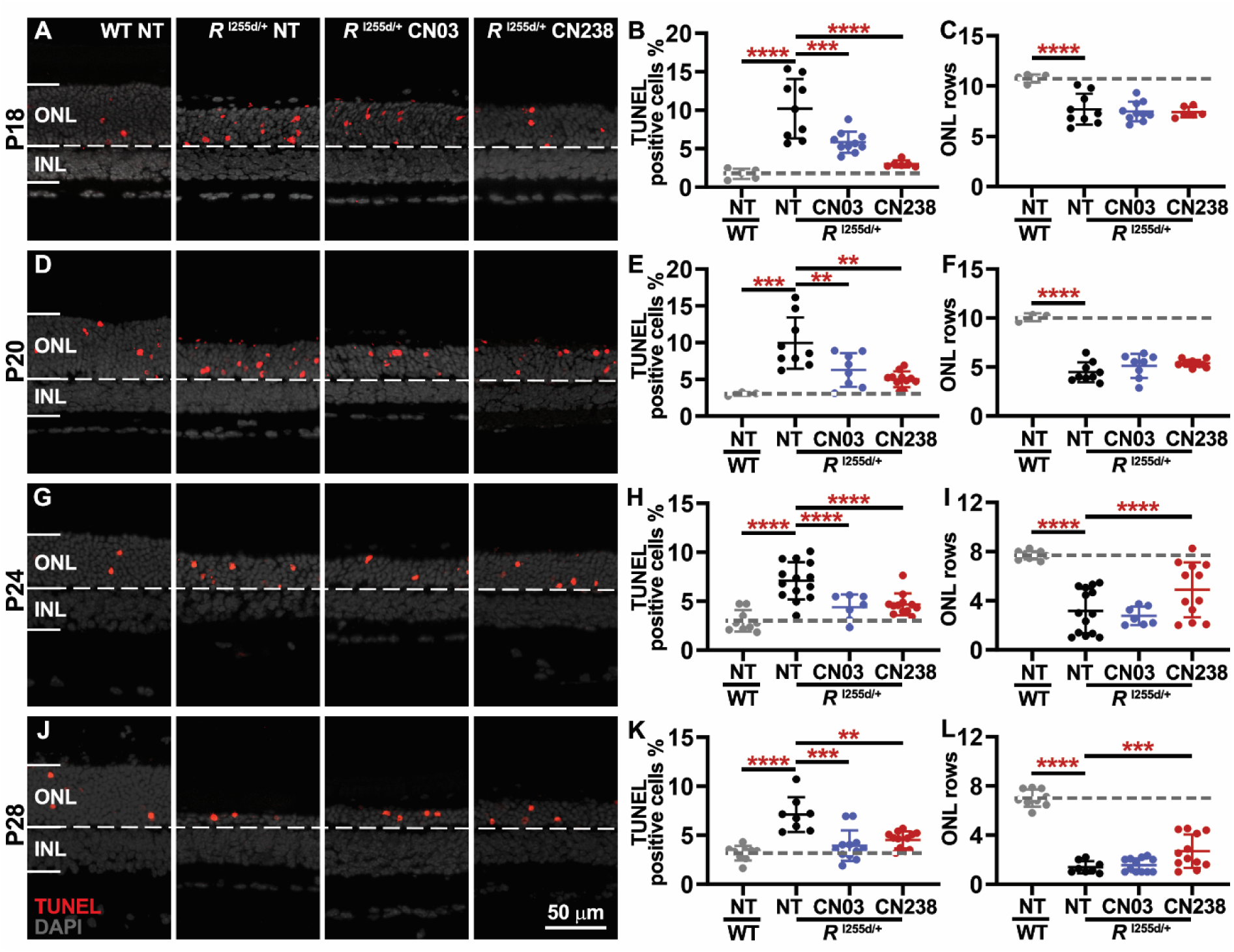
cGMP analogues preserve *Rho*^I255d/+^ photoreceptor viability. Organotypic retinal explant cultures were derived from wild-type (WT) and *Rho*^I255d/+^ mice at post-natal (P) day 12, cultured without treatment for the first two days, and then treated or not with 50 µM of CN03 or CN238 for further 4, 6, 10, or 14 days *in vitro*. (**A, D, G, J**) Representative sections of retinal explants cultured until P18 (**A**), P20 (**D**), P24 (**G**), and P28 (**J**) and stained with TUNEL assay (red). DAPI (grey) was used as nuclear counterstain. Non-treated (NT) WT and *Rho*^I255d/+^ cultures shown for comparison. (**B, E, H, K**) Percentages of TUNEL-positive, dying cells in the outer nuclear layer (ONL) of P18 (**B**), P20 (**E**), P24 (**H**), and P28 (**K**) retinas. NT mutant retina showed increased numbers of dying cells compared with WT, while treatment with CN03 and CN238 reduced TUNEL-positive cells. (**C, F, I, L**) Quantification of ONL rows in *Rho*^I255d/+^ retinal explants treated until P18 (**C**), P20 (**F**), P24 (**I**), and P28 (**L**). The number of ONL rows in NT mutant retinas was consistently lower than NT WT at corresponding time points. CN238 preserved ONL rows in retina cultured until P24 and P28. n = 3-17 retinas from different animals; error bars indicate SD; statistical testing: Two-way ANOVA with Dunnett’s multiple comparisons test; significance levels: ** p ≤ 0.01, *** p ≤ 0.001, **** p ≤ 0.0001; INL = inner nuclear layer; scale bar = 50 µm.

### cGMP analogues decrease calpain-2 activation

Calpain-2 activity is considered a short-lived event in the late phase of the cell death process (15). The correlation between retinal degeneration and elevated calpain-2 activation in *Rho*^I255d/+^ (*cf*. Fig. 2) led us to examine the effect of cGMP analogues on calpain-2. Compared to the WT situation, in short- and long-term retinal explant cultures, the immunostaining for activated calpain-2 was elevated in NT *Rho*^I255d/+^ retina (Fig. S1, Table 7). This phenomenon appeared to decrease with longer culture durations (Fig. S1B, D, F, H). The cGMP analogues CN03 and CN238 significantly reduced the numbers of ONL cells showing activated calpain-2 immunopositivity in short-term cultures lasting until P18 and P20. Yet, in longer-term cultures, cGMP analogues had no significant effect on calpain-2 activation. These findings suggest that calpain-2 may not be involved in the neuroprotection afforded by cGMP analogues. Alternatively, calpain-2 activation could be a transient event evident only in the early phases of *Rho*^I255d/+^ retinal degeneration.

### PKG inhibition preserves cone photoreceptor viability in *Rho*^I255d/+^ retina

Similar to the human situation, the mouse ONL harbors rod and cone photoreceptors with an overall approximate ratio of 97:3 (17, 18). The data obtained thus far suggested that CN03 and CN238 protected rod photoreceptors in *Rho*^I255d/+^ retina. To assess how cone photoreceptors were affected by these cGMP analogues, we labelled cones with the specific marker arrestin-3 (Arr3) and quantified the number of cones per 100 µm of retinal circumference.

From P18 to P28, the numbers of cone photoreceptors showed a progressive decrease in NT mutant retina, when compared to WT (Fig. 4, Table 6). Treatment with both cGMP analogues improved cone survival, an effect that was especially evident with CN238 at P28 (Fig. 4H). Overall, these findings illustrated a remarkable cone photoreceptor protection conferred by the cGMP analogue CN238 and in relative terms, in the time-frame studied here, the cone protective effect appeared to exceed that of rod photoreceptors.

**Figure 4.**
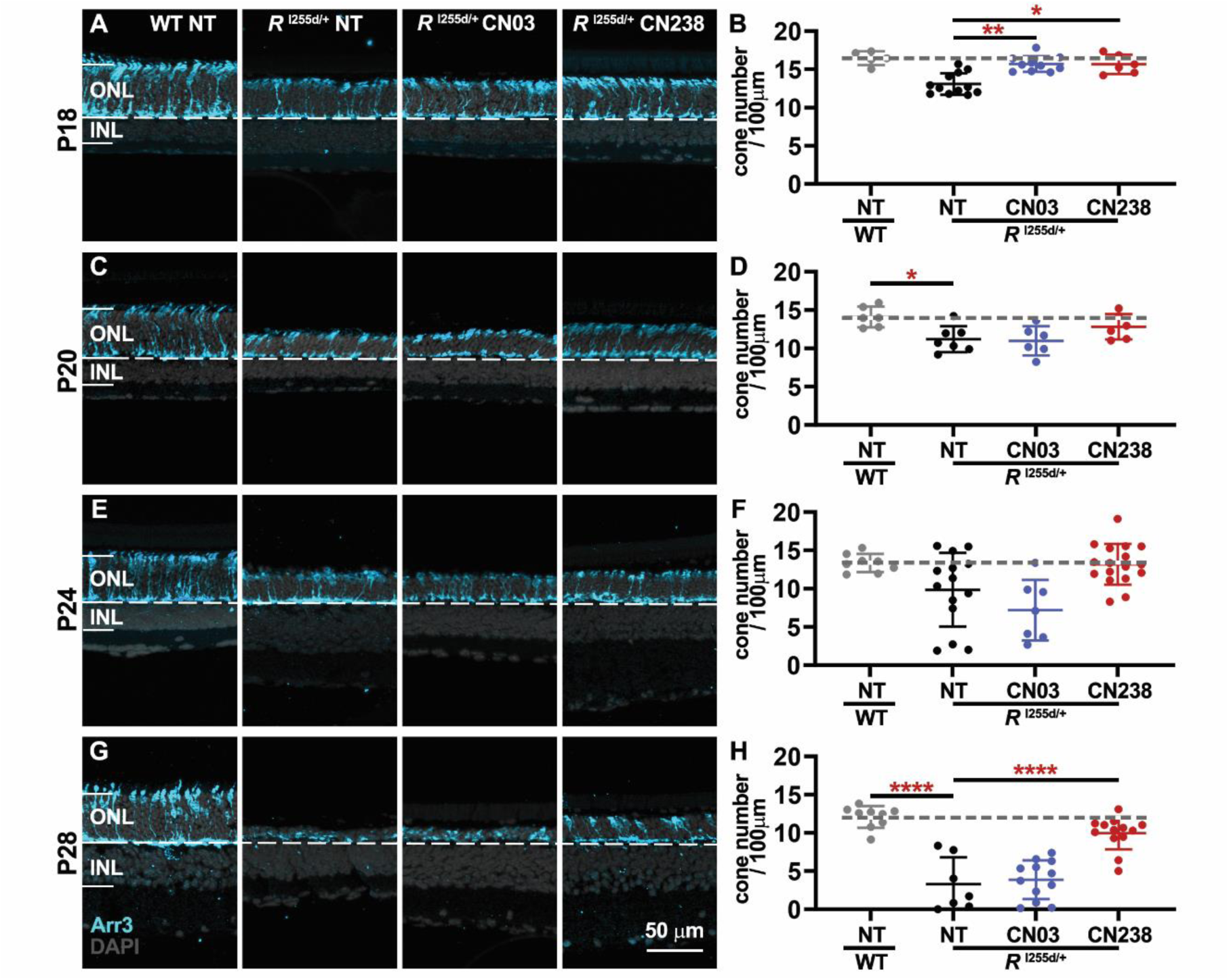
CN238 protects cone photoreceptors in the *Rho*^I255d/+^ retina. (**A, C, E, G**) In retinal explant cultures derived from wild type (WT) and *Rho*^I255d/+^ mice cone photoreceptors were labeled using cone arrestin-3 (blue; Arr3) immunolabelling. DAPI (grey) was used as nuclear counterstain. WT and *Rho*^I255d/+^ retinas were cultured from post-natal (P) day 12 to P18 (**A**), P20 (**C**), P24 (**E**), and P28 (**G**), treatment with either CN03 or CN238 was started at P14. (**B, D, F, H**) Quantification of cones per 100 µm of retinal circumference in P18 (**B**), P20 (**D**), P24 (**F**), and P28 (**H**) retina. Compared with non-treated (NT) WT, cone counts in NT mutant were decreased. In retinas cultured until P28, CN238 afforded significant cone protection, while CN03 had no effect. n = 5-17 retinas from different animals; error bars indicate SD; statistical testing: Two-way ANOVA with Dunnett’s multiple comparisons test; significance levels: * p ≤ 0.05, **** p ≤ 0.0001; INL = inner nuclear layer, ONL = outer nuclear layer; scale bar = 50 µm.

### cGMP analogues reduce non-apoptotic cell death markers but do not affect apoptosis

The enzymatic activities of calpain-type proteases, HDAC, and PARP were previously implicated in retinal degenerative mechanisms across various RP animal models, including the *Rho*^I255d/+^ mouse (4, 13). Additionally, poly (ADP-ribose) (PAR), *i.e.* the product of PARP activity, was shown to accumulate in degenerating photoreceptors (4, 19).

In retinal explant cultures at P20, an enzymatic *in situ* calpain activity assay revealed increased calpain activity in the NT *Rho*^I255d/+^ ONL compared with NT WT (Fig. S2A). Notably, both CN03 and CN238 reduced calpain activity significantly (Fig. S2B, Table 8). The HDAC *in situ* activity showed a trend towards elevation in NT mutant retina that was not significantly changed by the cGMP analogues CN03 and CN238 (Fig. S2C, D, Table 8). PARP activity and PAR accumulation (Fig. S2E-H, Table 8) were elevated in NT mutant compared with NT WT. Notably, CN238 treatment significantly reduced PARP activity and PAR accumulation, while CN03 treatments were not significant.

Apoptosis was previously suggested as one cell death mechanism contributing to the loss of *Rho*^I255d/+^ photoreceptors (13). Notably, cleaved, activated caspase-3, a protease critical for the execution of apoptosis, was significantly elevated in *Rho*^I255d/+^ retina at P20 (13). To investigate whether caspase-3 activation was altered by PKG inhibition, we quantified the numbers of ONL cells showing activated caspase-3 immunoreactivity in retinal explants cultivated until P20. While the numbers of ONL cells displaying caspase-3 activation were clearly elevated in the mutant retina (Fig. S2I, J, Table 8), overall, they were far lower than the number of cells labelled by the TUNEL assay (*cf*. Fig. 2). Treatment with either CN03 or CN238 had no significant effect on caspase-3 positive cell counts.

In summary, these findings indicated that calpain and PARP activity, as well as accumulation of PAR, were likely involved in excessive cGMP- and PKG-induced cell death. Conversely, both HDAC and caspase-3 activity seemed to be unrelated to (excessive) cGMP-signaling. Hence, cGMP analogues inhibiting PKG may only address a part of the cell death mechanisms driving photoreceptor loss in the *Rho*^I255d/+^ ADRP mouse model.

### CN238 treatment maintains photoreceptor light responsiveness

Our data thus far indicated a strong neuroprotective effect of the cGMP analogue CN238 on rod and cone photoreceptors in the *Rho*^I255d/+^ retina. To study whether and to what extent this protection translated into a functional benefit, we used micro electroretinogram (µERG) recordings performed on micro-electrode array (MEA) on P24 *in vitro* retina (20).

Acutely explanted *Rho*^I255d/+^ retina showed minimal µERG responses to photopic flashes of light, with negative deflection a-wave amplitudes (*i.e.* the primary responses of photoreceptors) remaining close to noise level (−7.44 ± 2.46; Fig. 5A). In *Rho*^I255d/+^ retina explanted at P12 and maintained in culture until P24 (*i.e.* 12 days *in vitro*) photoreceptor responses to light were almost entirely absent (−2.47 ± 2.23 µV). In stark contrast to this, *Rho*^I255d/+^ explants cultured for 12 days *in vitro* and treated with CN238 showed a pronounced 8.1-fold increase in light responses (−20.02 ± 12.49 µV; Fig. 5A, B). This increase was statistically significant towards both untreated retinal cultures (*p* = 0.0045) and towards acutely explanted retina (p = 0.039). As opposed to conventional ERG recordings the µERG technique using MEA allows an analysis of photoreceptor light responsiveness in the spatial domain (Fig. 5C). This analysis over 59 MEA electrodes revealed that the CN238 treatment effect is essentially uniform over the entire retinal surface.

**Figure 5.**
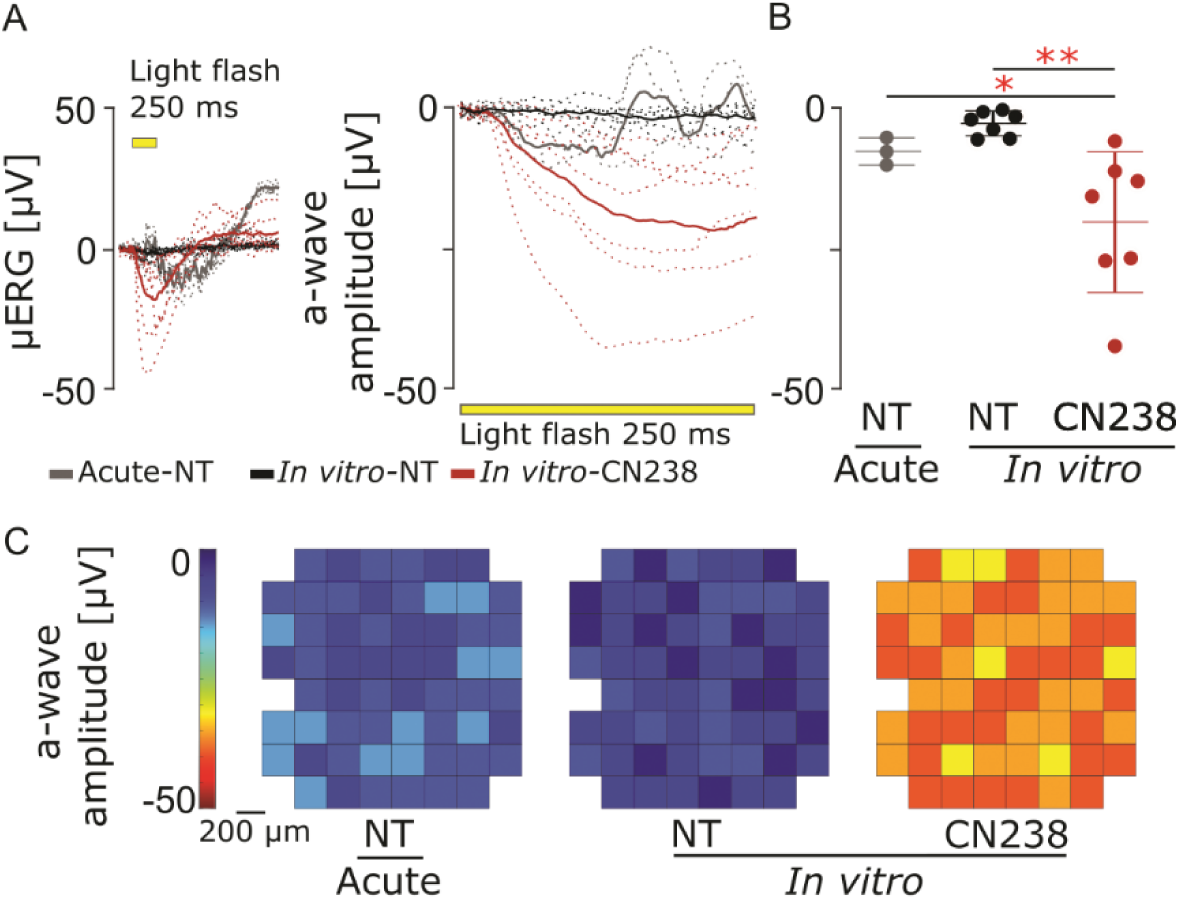
CN238 improves retinal light responsiveness. Assessment of retinal light responsiveness by micro-electroretinogram (µERG), using multi-electrode array (MEA). **A.** µERG traces of retinal explants; non-treated (NT), acute retinal explants from *Rho*^I255d/+^ were recorded at post-natal (P) day 24 (grey; dashed lines: single recordings, solid line: grand mean). Organotypic retinal explants were obtained from *Rho*^I255d/+^ at P12, treated with cGMP analogue CN238 from *Rho*^I255d/+^ retinas from P14 onwards, and recorded at P24 (NT in black and CN238 in red). Each retinal explant was stimulated by 5 consecutively applied full-field flashes (250 ms, yellow bar). **B**. A-wave peak amplitudes from data shown in A. **C.** Representative activity heatmap of best-performing condition (59 recordings MEA electrodes, 200 µm electrode interspace), indicating the treatment effect over the entire retinal surface (1.6 mm2). n = 3-7 retinas from different animals; error bars indicate SD; Statistical testing: One-way ANOVA with Dunnett’s multiple comparisons test; significance level: * p ≤ 0.05, ** p ≤ 0.01.

To conclude, treatment with the PKG inhibitor CN238 not only protected photoreceptor viability but also preserved light responsiveness.

## Discussion

Evidence of accumulated cGMP in photoreceptors has been found in various RP models, and has been implicated as a critical causative factor of photoreceptor degeneration (5). In the present study, we showed that excessive cGMP-signaling also contributes to photoreceptor degeneration in the *Rho*^I255d^ ADRP mouse model. Notably, we found that cGMP analogues inhibiting PKG provided long-lasting neuroprotection for both rod and cone photoreceptors.

### cGMP accumulation in the *Rho*^I255d^ ADRP model

cGMP is a secondary messenger, which in photoreceptors is synthesized by retinal guanylyl cyclase (retGC). cGMP plays a pivotal role in the vertebrate phototransduction cascade within the OS (21, 22), where its physiological concentration is thought to be in the 1-5 µM range (23). Excessive cGMP appears to be a causative factor that leads to photoreceptor cell death in a number of different RP animal models, including the *rd1*, *rd10*, and *R562W*\**V685M* ARRP mouse models as well as the *Rho*^P23H^ and *Rho*^S334ter^ ADRP rat models. In these models, the mutated gene would normally be expressed in the rod OS (4, 14, 24, 25). cGMP accumulation is observed mostly within the OS/IS in *Rho*^P23H^ and *Rho*^S334ter^ rats, but may extend to the entire photoreceptor cell in *rd1*, *rd10*, and *R562W*\**V685M* mice (4, 14, 25). In contrast, cGMP specifically accumulated in photoreceptor segments of *Rho*^I255d/+^ and *Rho*^I255d/I255d^ mice.

The mechanisms underlying these phenotype-specific differences remain unclear but may involve disrupted ciliary transport or alterations in cGMP synthesis. Indeed, specific rod deletion of ITF172, a component of the connecting cilium, results in RP and is associated with rod OS protein mis-localization, such as rhodopsin (26). In line with this, rhodopsin mis-localization was observed during retinal degeneration in homozygous *Rho*^I255d/I255d^ and heterozygous *Rho*^I255d/+^ mice (13). Thus, there is the possibility that an impaired intraflagellar transport caused by the *Rho*^I255d^ mutation also results in improper distribution of either cGMP itself or the cGMP synthesis or degradation machinery.

### PKG inhibition delays photoreceptor loss in *Rho*^I255d/+^ retina

The first- and second-generation selective PKG inhibitors CN03 and CN238, both analogues of cGMP, were previously shown to preserve photoreceptors *in vitro* and *in vivo* in the *rd1*, *rd2*, and *rd10* mouse models for ARRP (10, 11, 27). Here, we demonstrated that both cGMP analogues partially counteracted the progression of *Rho*^I255d^ mutation-induced ADRP. The rate of ONL cell death, as evidenced by the TUNEL assay, was reduced in both short- and long-term retinal explant cultures, with CN238 showing the stronger photoreceptor neuroprotection.

Importantly, CN238 treatment also rescued retinal function as assessed by µERG recordings (20), highlighting the compound’s therapeutic potential. Since the light intensity for retinal stimulation used in these experiments was in the photopic range, this result pointed at a significant functional protection of the retinás cone photoreceptors.

### Pro-survival effect of CN238 on cone photoreceptors

In RP, the viability of cone photoreceptors is compromised secondarily, once most rod photoreceptors have been lost to the disease. The mechanisms behind this secondary cone loss are still largely unclear even though a general therapeutic concept for their preservation is to “spare rods to save the cones” (28). The compound CN238 was previously shown to preserve cones in the *rd10* mouse model for ARRP (11), and here we confirmed a marked cone neuroprotective effect in the *Rho*^I255d/+^ model for ADRP, for treatment periods lasting up to 14 days. At present it is unclear whether this cone protection is indirect, afforded by the preservation of rods, or whether CN238 had a direct effect on cone viability.

From studies on the CNG-channel deficient *Cnga3*^-/-^ cone degeneration mouse model, we know that increased PKG activity can cause cone death (29). Conversely, in the *Cnga3*^-/-^ model the *in vivo* inhibition of PKG, with either the cGMP analogue (Rp)-8-Br-cGMPS or the ATP-site inhibitor KT5823, prevented cone degeneration (30). From this data it thus seems possible that CN238 may have had a direct cone protective effect also in *Rho*^I255d/+^ retina.

Conversely, in a rod dominated retina CN238 may have preserved cones indirectly, via the promotion of rod survival. Recent evidence suggests that rods and cones use different metabolic pathways to satisfy their high energy demands (31, 32), and cones may be metabolically dependent on rods (33). Hence, it seems likely that a preservation of rods via CN238 treatment contributed to the cone preservation observed. Future metabolomic studies, including metabolic tracing, may be used to investigate to what extent PKG inhibitors act on rods and/or cones.

### Apoptotic and non-apoptotic cell death of *Rho*^I255d/+^ photoreceptors

Cell death in *Rho*^I255d/+^ retina can be executed by both apoptosis and non-apoptotic cGMP-dependent cell death (5, 13) as shown by increased cellular activities of, on the one hand, caspase-3 activation and, on the other hand, calpain- and PARP activity. In the present study, we found that PKG inhibitors preserving *Rho*^I255d/+^ photoreceptors also reduced the activity of calpain and PARP. However, at the same time, these PKG inhibitors had no significant effect on HDAC activity and caspase-3 activation. This outcome suggests that apoptotic photoreceptor cell death is not impeded by PKG inhibition. While this may somewhat reduce the therapeutic efficacy of PKG inhibition in ADRP, it may also reduce detrimental side effects, notably because an indiscriminate inhibition of apoptosis could favor cancerogenesis (34).

The effects of PKG inhibitors on calpain activity may be particularly interesting. Theoretically, these could be due to direct inhibition of CNG-channels in photoreceptor OS. However, cGMP analogues are known to inhibit CNG-channels with an efficacy around 2 log units lower compared to PKG (35), making direct effects on the channel less likely. In view of recent data on the involvement of T-type voltage-gated Ca^2+^-channel (VGCC) and the Na^+^/Ca^2+^-exchanger (NCK) in photoreceptor degeneration in the *rd1* mouse model for ARRP, it is plausible to think that calpain becomes activated by processes further down-stream of elevated cGMP-levels (16).

When the relative numbers of cells positive for calpain activity and activated caspase-3 are compared to the numbers of dying cells, it seems that calpain activity, and by inference cGMP-dependent cell death, is more important for *Rho*^I255d/+^ photoreceptor degeneration. Curiously, HDAC activity, formerly clearly linked to cGMP-dependent cell death (36, 37), in the *Rho*^I255d/+^ model for ADRP appears to be independent of this pathway. Overall, these findings highlight the diversity of cell death mechanisms triggered by the *Rho*^I255d^ mutation.

## Conclusion

This study focused on a recently generated mouse model carrying a single copy of a mutant *RHO* gene, mimicking the homologous human disease ADRP. We found excessive cGMP-signaling and PKG activity to be a driver of photoreceptor degeneration in *Rho*^I255d/+^ retina, providing for a common therapeutic target shared across dominant and recessive RP disease models. This expands the therapeutic scope of cGMP analogues such as CN238, bringing us closer to addressing the unmet needs of RP patients in ways that are both effective and economical. Further studies aimed to promote clinical feasibility will focus on establishing a suitable administration paradigm and drug delivery approach to allow cGMP analogues to reach retinal photoreceptors.

## Methods

### Animals

C57BL/6J wild-type (WT) and *Rho*^I255del^ mutant mice (*Rho*^I255d^) were used for this project (13). WT animals were crossed with homozygous *Rho*^I255d/I255d^ mice to generate heterozygous *Rho*^I255d/+^ animals. All efforts were made to minimize the number of animals used and their suffering. Animals were taken care of under standard white cyclic lighting, had free access to water and food, and were used irrespective of gender. Protocols compliant with the Association for Research in Vision and Ophthalmology (ARVO) statement for the use of animals in vision research and the German law on animal protection were reviewed and approved by the "Einrichtung fuer Tierschutz, Tierärztlichen Dienst und Labortierkunde" of the University of Tuebingen (Registration Nos: AK02-20M).

### Organotypic retinal explants

The retinal explanation procedure was followed by a protocol previously described (38). Briefly, WT and heterozygous *Rho*^I255d/+^ were euthanized at post-natal (P) day 12 through CO_2_ inhalation. The eyes were enucleated and placed 5 min at room temperature (RT) into defined R16 basal medium (07491252A, BM, Gibco, Paisley, UK), then incubated 15 min at 37 °C in BM with 0.12% proteinase K (39450, MP Biomedicals, Irvine, CA, USA). Proteinase K activity was blocked by 20% fetal bovine serum (F7524, FBS, Sigma, Hamburg, Germany) for 5 min at room temperature (RT). The anterior segments were carefully removed from the eyeballs, the optic nerve was cut, and the RPE-attached retinas were removed from the sclera. Retinas were cut in a clover leaf-like shape and transferred to a culturing membrane insert in a six-well plate (Corning Life Sciences, Corning, NY, USA) with the RPE facing down. Subsequently, it was incubated with 1.2 mL R16 complete medium (CM) with supplements, free of serum and antibiotics. Explants were incubated in a humidified incubator with 5% CO_2_ at 37 °C. After 48 h, the retina was either exposed to 50 µM (10, 11) of CN03 or CN238 (Biolog Life Science Institute GmbH, Bremen, Germany), dissolved in water, or kept as an untreated control. The medium was changed every second day and cultures were ended at 4 and 6 days of treatment (short-term cultures, ending at P18 or P20, respectively), or at 12 and 14 days of treatment (long-term cultures, ending at P24 or P28, respectively). For MEA-recording procedures, retinal dissections and medium-changes were performed in the dark under red light, treatments with 50 µM CN238 started at P14 and ended at P24.

## Histology

### Fixed and unfixed sections

For retinal cross-section preparation, WT, *Rho*^I255d/+^, and *Rho*^I255d/I255d^ mice were enucleated at P20 and fixed in 4% paraformaldehyde (PFA) for 45 mins. Afterwards, the anterior segment, lens, and vitreous body was removed. Retinal explants, after culturing, were fixed directly with 4% PFA. The following steps were common for both preparations: tissue sections were incubated in graded solutions containing 10%, 20%, and 30% sucrose and then embedded in Tissue-Tek O.C.T. compound (Sakura Finetek Europe, Alphen aan den Rijn, Netherlands) and snap-frozen on liquid N_2_. Unfixed sections (i.e. for calpain, PARP activity assays) were embedded in Tissue-Tek O.C.T. compound and directly frozen in liquid N_2_. Tissue sections of 14 µm were prepared using a NX50 microtome (Thermo Scientific, Waltham, MA, USA) and thaw-mounted onto Superfrost Plus glass slides (R. Langenbrinck, Emmendingen, Germany), then stored at −20 °C.

### TUNEL assay

Cell death was detected by terminal deoxynucleotidyl transferase dUTP nick end labeling (TUNEL) assay (11684795910, Sigma-Aldrich *in situ* Cell Death Detection Kit, red fluorescence). Fixed sections were dried at 37°C for 30 min, then washed in phosphate-buffered saline (PBS) solution at RT, and unfixed sections were fixed by 4% PFA for 10 min in parallel, then washed with PBS. Both fixed and unfixed tissues shared the same steps afterward. Nucleases were inactivated by placing the sections in Tris buffered saline (TBS) with proteinase K at 37 °C for 5 min. Sections were then washed for 3 times in TBS for 5 min. Subsequently, sections were placed in ethanol-acetic acid mixture (70:30) at −20°C for 5 min, followed by 3 washes in TBS and incubation in blocking solution (BS; 10% normal goat serum, 1% bovine serum albumin, 1% fish gelatin in 0.1% PBS-Triton X-100) for 1 h at RT. Slides were incubated in the TUNEL reaction solution at 4°C overnight, and then washed 2 times in PBS. Finally, the tissue sections were mounted with Vectashield with 4′,6-diamidino-2-phenylindole (DAPI; Vector Laboratories, Newark, CA, USA), here DAPI served as nuclear counterstain.

### Immunofluorescence

Fixed slides were placed in BS for 1 h at RT. Primary antibodies against sheep cGMP (1: 100, Prof. Harry Steinbusch, Maastricht University, Netherlands), rabbit calpain-2 (1:200, ab39165, Abcam, Cambridge, UK), rabbit cone arrestin-3 (1:500, ab15282; Millipore, USA), rabbit cleaved caspase-3 (1:100, #5A1E, cell signaling, Danvers, USA) were diluted in the BS and incubated at 4 °C overnight. Subsequently, secondary antibodies conjugated with Alexa Fluor 488 or 568 (Thermo Fisher Scientific, Sindelfingen, Germany) were applied to slides for 1 h at RT. Lastly, slides were washed with PBS and mounted with DAPI (Vector Laboratories, Newark, CA, USA).

### *In situ* activity assays

#### Calpain activity

This assay allows detecting the overall calpain activity *in situ* on unfixed tissue sections, following a protocol described previously (15). Unfixed sections were rehydrated in calpain reaction buffer (CRB; 5.96 g HEPES, 4.85 g KCl, 0.47 g MgCl_2_, and 0.22 g CaCl_2_ in 100 mL ddH_2_O; pH 7.2) for 15 min at RT. Sections were then placed in CRB with t-BOC-Leu-Met-CMAC (25 µM; A6520, Thermo Fisher Scientific) with 2 mM dithiothreitol (DTT), incubated for 3.5 h at 37 °C in the dark, and subsequently incubated for 30 min at RT with ToPro3 (1:1 000 in PBS, T3605, Invitrogen, Carlsbad, USA), which was used as the nuclear counterstain. Afterward, sections were mounted using Vectashield without DAPI (Vector Laboratories, Newark, CA, USA) for immediate visualization.

#### PARP activity

This assay allows testing the overall activity of PARP on unfixed tissue sections (39). Sections were rehydrated with Tris buffer, then incubated with fluorescent NAD^+^ and the reaction mixture (1 mM DTT, 50 μM 6-Fluo-10-NAD^+^; Biolog, Cat. Nr.: N 023), and PARP buffer (10 mM MgCl_2_ in 100 mM Tris buffer with 0.2% Triton X100) for 3.5 h at 37 °C. Then, sections were mounted in Vectashield with DAPI (Vector Laboratories, Newark, CA, USA) for immediate visualization.

#### HDAC activity

The assay allows to resolve the overall HDAC activity on fixed sections. Sections were incubated in the assay buffer for 30 min, then they were exposed to Fluor de lysate Sirt 2 substrate working solution 100 µL (3 µL FLUOR DE LYS® -SIRT2 deacetylase substrate (200 µM; Enzo Life Sciences, New York, USA) + 4 µL NAD^+^ (1mM) + 1 µL NP-40 (0.1%) + 92 µL assay buffer) at 37 °C for 3 h. Afterwards, the slides were fixed in methanol at −20 °C for 20 min. After staining with ToPro3 (1:1000, T3605, Invitrogen, Carlsbad, USA) for 30 min, sections were mounted with FLUOR DE LYS® developer II concentrate (BML-KI176-1250, Enzo Life Sciences, New York, USA), incubated overnight, and imaged the following day.

#### PAR-DAB staining assay

To investigate PAR accumulation in the retina, DAB (3,3′-Diaminobenzidine) staining (39) was performed on fixed sections. Sections were incubated with quenching of endogenous peroxidase activity (40% MeOH and 10% H_2_O_2_ in 0.3% PBST) for 20 min, followed by incubation in 10% NGS, 1% BSA, and 0.3% PBST for 30 min at RT, then incubated with primary antibody against mouse PAR (1:200; ALX-804-220-R100; Enzo Life Sciences, Farmingdale, NY, USA) overnight at 4°C. After 1 h incubation with Vector ABC Kit (Vector Laboratories, solution A and solution B in PBS, 1:150 each), followed with biotinylated secondary antibody (1:150, Vector in 5% NGS in PBST) for 1 h at RT, DAB staining solution (0.05 mg/mL NH 4 Cl, 200 mg/mL glucose, 0.8 mg/mL nickel ammonium sulphate, 1 mg/mL DAB, and 0.1 vol. % glucose oxidase in phosphate buffer) was applied immediately, incubated for 3 min 10 s, and then rinsed with phosphate buffer to stop the reaction. Slides were mounted in Aquatex (Merck, Darmstadt, Germany).

#### Microscopy and image processing

A Zeiss Imager Z.2 fluorescence microscope, equipped with an ApoTome 2, an Axiocam 506 mono camera, and HXP-120V fluorescent lamp (Carl Zeiss Microscopy, Oberkochen, Germany) was used for microscopy. The excitation (λExc.)/emission (λEm.) characteristics of the filter sets used for the different fluorophores were as follows (in nm): DAPI (λExc. = 369 nm, λEm. = 465 nm), AF488 (λExc. = 490 nm, λEm. = 525 nm), and ToPro3 (λExc. = 642 nm, λEm. = 661 nm). Images were captured using a 20x/0.8 objective and Zen 2.3 blue edition software. Sections of 14 µm thickness were analyzed using 9 consecutive images using the Z-stack function. Adobe Photoshop 2020 (San Jose, CA, USA) was used for image processing and assembly of final figures for publication.

#### Quantification

Images were captured on 3 sagittal retinal sections from at least 3 different animals per genotype. The average cell size occupied by a photoreceptor cell was determined by counting DAPI- or ToPro3-stained nuclei in nine different areas of ONL. The number of positively labeled cells (*i.e.* TUNEL, caspase-3, PARP, calpain, and calpain-2) in the ONL was manually counted. The percentage of positive cells was calculated from the ratio of the number of positive cells divided by the total number of ONL cells (ONL area divided by the average cell size). Arrestin-3-labeled cells in the ONL were counted manually and their numbers expressed as cells per 100 μm of retinal circumference.

#### Statistics

Error in graphs and text is given as standard deviation (SD). Values are given as mean ± SD. GraphPad Prism 10.1.2 software (GraphPad Software, CA, USA) was used for statistical analysis, unpaired Student’s *t*-test was used for 1:1 comparisons, one-way ANOVA and Two-way ANOVA with Dunnett’s multiple comparisons test were performed to compare more than two groups. Levels of significance were: * *p* ≤ 0.05, ** *p* ≤ 0.01, *** *p* ≤ 0.001, **** *p* ≤ 0.0001.

### *Ex vivo* retinal function test

#### Micro-electroretinogram (µERG)

To estimate the retinal light responsiveness, the method of micro-electroretinography (20) was utilized. Retinal explants were kept dark-adapted, and handling procedures, including dissection and medium change, were carried out in the dark under dim red-light conditions. For recordings, a micro-electrode array system (MEA; USB-MEA60-Up-BC-System-E, Multi-Channel Systems; MCS; Reutlingen, Germany), equipped with a 60MEA 200/30iR-ITO-pr (59 recording electrodes, 1 internal reference electrode) was utilized and unfiltered raw data were collected at a sampling rate of 25 kHz. In experiments, retinal explants were placed centrally, ganglion cells facing down, on the electrode field in the recording chamber filled with drug-free R16 CM at 37°C, and were kept in a carbonated environment throughout the whole recording. The light stimulation was applied using a white light source (LED, 2350 mW, MCWHD3, Thorlabs, Bergkirchen, Germany) and guided beneath the transparent glass MEA using fiber-optics. The operation of the light simulation (LEDD1B T-Cube, Thorlabs) was programmed using MATLAB (2019b, Mathworks, Natick, MA, USA) and trigger synchronized with the MEA-recordings controlled by a dedicated protocol implemented within the MC_Rack software (v4.6.2, MCS) and the digital I/O-box (MCS, Reutlingen, Germany). A spectrometer USB4000-UV-VIS-ES (Ocean Optics, Ostfildern, Germany) was used to calibrate the intensity of the applied light flashes (4.20 × 10 ^13^ photons/cm^2^/s). Five full-field flashes of 250 ms duration in a 20 s interval were applied per recording.

#### Data processing and evaluation

The recorded MEA files were filtered using the Butterworth 2^nd^-order (MC_Rack, v4.6.2, MCS) to extract field potentials (band pass 2-40 Hz). They were then converted to *.hdf files using the MC DataManager (v1.6.1.0, MCS, Reutlingen, Germany) and imported into MATLAB using the McsMatlabDataTools software bundle (MCS, Reutlingen, Germany). To obtain the light-evoked photoreceptor response, the light-correlated maximum negative deflection, the a-wave, was extracted (20).

## Supporting information

supplemental results

## Acknowledgments

The authors would like to thank Norman Rieger (Institute for Ophthalmic Research, Eberhard-Karls-Universität Tübingen) for excellent technical assistance.

## Author contributions

Conceptualization, Y.Z. and F.P-D.; methodology, Y.Z., L.P., P.N-K. and W.H.; software, Y.Z. and W.H.; validation, Y.Z.; formal analysis, Y.Z. and W.H.; investigation, Y.Z., L.P. and P.N-K.; data curation, Y.Z.; writing-original draft preparation, Y.Z., L.P, and W.H.; writing—review and editing, C.H. and F.P-D.; supervision, F.P-D.; project administration, F.P-D.; funding acquisition, F.P-D. All authors have read and agreed to the published version of the manuscript.

## Data Availability Statement

All data generated or analyzed during this study are included in this published article and its Supplementary Materials Files.

