## supplemental results for "Protein kinase G inhibition preserves photoreceptor viability and function in a new mouse model for autosomal dominant retinitis pigmentosa"

**Supplementary results**

**Figures**

Fig. S1. cGMP analogues decrease calpain-2 activation in short-term culture.

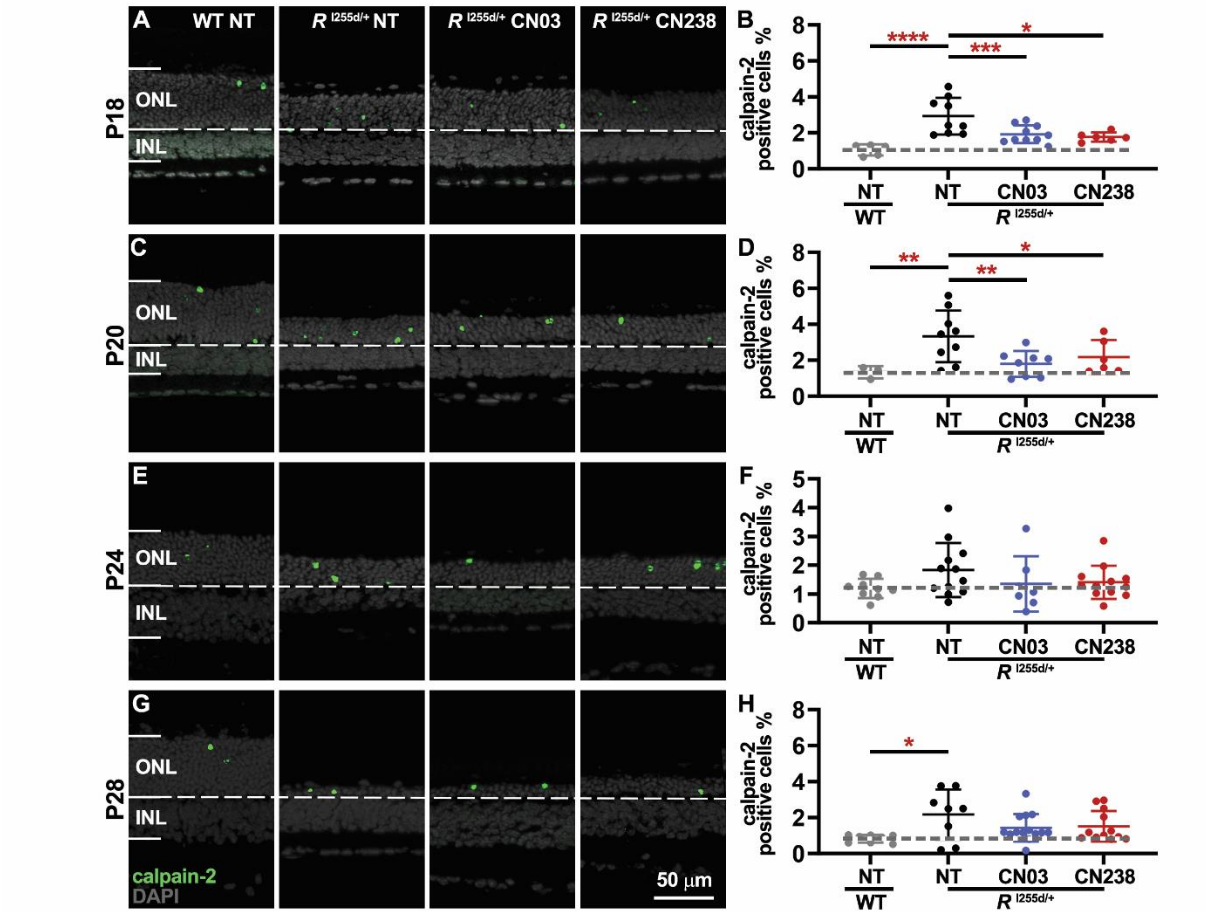


**Figure S1. cGMP analogues can reduce calpain-2 activation in *Rho*$^{I255d/+}$ retina.** (**A, C, E, G**) Organotypic retinal explant cultures were derived from wild-type (WT) and *Rho*$^{I255d/+}$ mice at post-natal (P) day 12, cultured without treatment for the first two days, and then treated or not with 50 µM of CN03 or CN238 for further 4, 6, 10, or 14 days *in vitro*. (**A, C, E, G**) Representative sections of retinal explants cultured until P18 (**A**), P20 (**C**), P24 (**E**), and P28 (**G**) and stained with an antibody directed against activated calpain-2 (green). DAPI (grey) was used as nuclear counterstain. Non-treated (NT) WT and *Rho*$^{I255d/+}$ cultures shown for comparison. (**B, D, F, H**) Quantification of calpain-2-positive cells in the outer nuclear layer (ONL) at P18 (**B**), P20 (**D**), P24 (**F**), and P28 (**H**). NT mutant exhibited an increased percentage of calpain-2-positive cells compared with NT WT at all time points except P24. cGMP analogues reduced calpain-2 activation in short-term cultures until P20. n = 3-14 retinas from different animals; error bars indicate SD; statistical testing: Two-way ANOVA with Dunnett's multiple comparisons test; significance level: * $p \leq 0.05$, ** $p \leq 0.01$, *** $p \leq 0.001$, **** $p \leq 0.0001$; INL = inner nuclear layer; scale bar = 50 µm.

Fig. S2. cGMP analogues reduce non-apoptotic photoreceptor cell death without decreasing apoptosis.

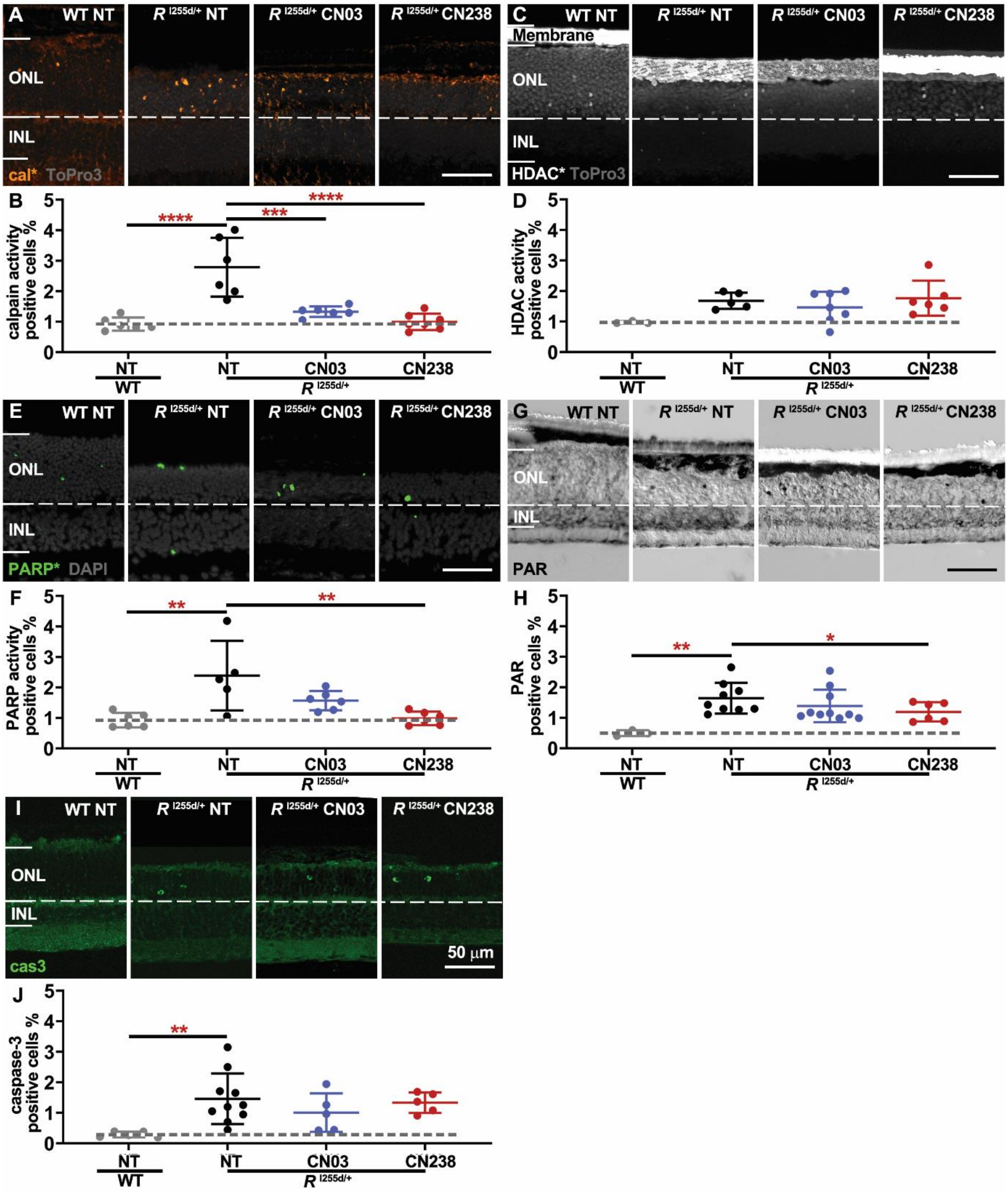


**Figure S2. cGMP analogues reduce photoreceptor cell death without decreasing apoptosis.** Organotypic retinal explants were obtained from *Rho*$^{I255d/+}$ and wild-type (WT) mice at post-natal (P) day 12 and were treated with cGMP analogues from P14 to P20. **A.** Staining for *in situ* calpain activity (orange). **B.** Percentage of calpain-activity positive cells in the outer nuclear layer (ONL). In *Rho*$^{I255d/+}$, CN03 and CN238 decreased the percentage of calpain-activity-positive cells compared to the non-treated (NT) situation. **C.** Staining for *in situ* HDAC activity (white). **D.** Quantification of histone deacetylase (HDAC) -activity-positive cells in the ONL showed no significant difference between NT and treatments. **E.** Staining for *in situ* poly (ADP-ribose) polymerase (PARP) activity (green). **F.** Quantification of PARP-activity-positive cells in the ONL revealed an increased percentage in NT *Rho*$^{I255d/+}$ compared to WT. Treatment with CN238 significantly decreased PARP activity in *Rho*$^{I255d/+}$ mutant retina, whereas CN03 had no effect. **G.** Staining for poly (ADP-ribose) (PAR; black). **H.** Quantification of PAR-labeled cells showed an increase in mutant NT retinas compared to WT. CN238 decreased the number of PAR-positive cells in *Rho*$^{I255d/+}$ mutant retina. **I.** Immunostaining (green) for cleaved caspase-3 (cas3). **J.** The percentage of caspase-3-positive cells was elevated in NT mutant compared with NT WT. cGMP analogues had no effect on the numbers of ONL caspase-3 positive cells. DAPI and ToPro3 (both in grey) were used for nuclear counterstaining. n = 3-10 retinas from different animals; error bars indicate SD; statistical testing: Two-way ANOVA with Dunnett's multiple comparisons test; significance level: * $p \leq 0.05$, ** $p \leq 0.01$, *** $p \leq 0.001$, **** $p \leq 0.0001$; INL = inner nuclear layer; scale bar = 50 µm.

### Tables

**Table 1.** cGMP accumulation in segments of *Rho*$^{I255d/+}$ and *Rho*$^{I255d/I255d}$: Quantitative data for graph presented in Figure 1B.

| Genotype | Peak value (AU) | n |
|---|---|---|
| *R*$^{I255d/I255d}$ | 1174.93 | 3 |
| *R*$^{I255d/+}$ | 585.67 | 3 |
| WT | 237.87 | 3 |
| *R*$^{I255d/+}$ neg | 183.38 | 3 |

**Table 2.** Comparison of cGMP accumulation in inner and outer segments (IS/OS) and outer nuclear layer (ONL) of *Rho*$^{I255d/+}$ and *Rho*$^{I255d/I255d}$: Quantitative data for graphs presented in Figures 1C and 1D.

| Figure | Genotype | Mean ± SD (%) | *p* - value (compared with WT) | n |
|---|---|---|---|---|
| Fig. 1C | *R*$^{I255d/+}$ neg | 3.74 ± 0.34 | $p = 0.3464$ | 3 |
| | WT | 3.20 ± 0.16 | | 3 |
| | *R*$^{I255d/+}$ | 4.49 ± 0.54 | $p = 0.0153$ | 3 |
| | *R*$^{I255d/I255d}$ | 5.39 ± 0.55 | $p = 0.0006$ | 3 |
| Fig. 1D | *R*$^{I255d/+}$ neg | 51.44 ± 7.19 | $p = 0.1503$ | 3 |
| | WT | 142.36 ± 41.26 | | 3 |
| | *R*$^{I255d/+}$ | 189.70 ± 64.47 | $p = 0.5746$ | 3 |
| | *R*$^{I255d/I255d}$ | 387.61 ± 70.48 | $p = 0.0011$ | 3 |

**Table 3.** Elevated calpain-2 activation (calpain-2 staining) related to photoreceptor cell death (TUNEL assay) in *Rho*$^{I255d/+}$ retina: Quantitative data for graphs presented in Figures 2B and 2D.

| Figure | Genotype | Mean ± SD (%) | *p* - value (compared with WT) | n |
|---|---|---|---|---|
| Fig. 2B | WT | 0.18 ± 0.06 | | 3 |
| | *R*$^{I255d/+}$ | 8.38 ± 1.98 | $p = 0.0020$ | 3 |
| Fig. 2D | WT | 0.04 ± 0.01 | | 3 |
| | *R*$^{I255d/+}$ | 3.56 ± 1.24 | $p = 0.0079$ | 3 |

**Table 4.** Effect of cGMP analogues on photoreceptor cell death (TUNEL assay) in treatments lasting 4-14 days: Quantitative data for graphs presented in Figures 3B, E, H, K.

| Figure | Culture scheme | Genotype - treatment | Mean ± SD (%) | *p* - value (compared with $R^{I255d/+}$ - NT) | n |
|---|---|---|---|---|---|
| Fig. 3B | P12 - 18 | WT - NT | 1.73 ± 0.67 | $p < 0.0001$ | 5 |
| | | $R^{I255d/+}$ - NT | 10.21 ± 3.87 | | 9 |
| | | $R^{I255d/+}$ - CN03 | 5.85 ± 1.38 | $p = 0.0009$ | 10 |
| | | $R^{I255d/+}$ - CN238 | 3.03 ± 0.45 | $p < 0.0001$ | 6 |
| Fig. 3E | P12 - 20 | WT - NT | 3.04 ± 0.26 | $p = 0.0010$ | 3 |
| | | $R^{I255d/+}$ - NT | 9.94 ± 3.48 | | 9 |
| | | $R^{I255d/+}$ - CN03 | 6.29 ± 2.29 | $p = 0.0025$ | 8 |
| | | $R^{I255d/+}$ - CN238 | 5.02 ± 1.07 | $p = 0.0002$ | 11 |
| Fig. 3H | P12 - 24 | WT - NT | 3.00 ± 1.10 | $p < 0.0001$ | 9 |
| | | $R^{I255d/+}$ - NT | 7.14 ± 1.79 | | 17 |
| | | $R^{I255d/+}$ - CN03 | 4.40 ± 1.29 | $p < 0.0001$ | 7 |
| | | $R^{I255d/+}$ - CN238 | 4.68 ± 1.08 | $p < 0.0001$ | 17 |
| Fig. 3K | P12 - 28 | WT - NT | 3.17 ± 0.74 | $p < 0.0001$ | 9 |
| | | $R^{I255d/+}$ - NT | 7.11 ± 1.78 | | 8 |
| | | $R^{I255d/+}$ - CN03 | 3.93 ± 1.56 | $p = 0.0001$ | 12 |
| | | $R^{I255d/+}$ - CN238 | 4.53 ± 0.86 | $p = 0.0012$ | 12 |

**Table 5.** Effect of cGMP analogues on photoreceptor viability (ONL row counts) in treatments lasting 4-14 days: Quantitative data for graphs presented in Figures 3C, F, I, L.

| Figure | Culture scheme | Genotype - treatment | Mean ± SD (%) | *p* - value (compared with $R^{I255d/+}$ - NT) | n |
|---|---|---|---|---|---|
| Fig. 3C | P12 - 18 | WT - NT | 10.76 ± 0.41 | $p = 0.0003$ | 5 |
| | | $R^{I255d/+}$ - NT | 7.70 ± 1.53 | | 9 |
| | | $R^{I255d/+}$ - CN03 | 7.47 ± 0.97 | $p = 0.9941$ | 10 |
| | | $R^{I255d/+}$ - CN238 | 7.41 ± 0.53 | $p = 0.2138$ | 6 |
| Fig. 3F | P12 - 20 | WT - NT | 10.07 ± 0.41 | $p < 0.0001$ | 3 |
| | | $R^{I255d/+}$ - NT | 4.47 ± 1.01 | | 9 |
| | | $R^{I255d/+}$ - CN03 | 5.12 ± 1.23 | $p = 0.4157$ | 8 |
| | | $R^{I255d/+}$ - CN238 | 5.38 ± 0.35 | $p = 0.1540$ | 11 |
| Fig. 3I | P12 - 24 | WT - NT | 7.70 ± 0.34 | $p < 0.0001$ | 9 |
| | | $R^{I255d/+}$ - NT | 3.66 ± 1.97 | | 17 |
| | | $R^{I255d/+}$ - CN03 | 2.77 ± 0.74 | $p = 0.1390$ | 7 |
| | | $R^{I255d/+}$ - CN238 | 5.45 ± 2.08 | $p < 0.0001$ | 17 |
| Fig. 3L | P12 - 28 | WT - NT | 7.02 ± 0.70 | $p < 0.0001$ | 9 |
| | | $R^{I255d/+}$ - NT | 1.39 ± 0.49 | | 8 |
| | | $R^{I255d/+}$ - CN03 | 1.55 ± 0.55 | $p = 0.6110$ | 12 |
| | | $R^{I255d/+}$ - CN238 | 2.69 ± 1.35 | $p = 0.0005$ | 12 |

**Table 6.** Effect of cGMP analogues on cone photoreceptor survival (cone arrestin-3 staining) in treatments lasting 4-14 days: Quantitative data for graphs presented in Figures 4B, D, F, H.

| Figure | Cultured schema | Genotype - treatment | Mean ± SD (%) | *p* - value (compared with $R^{\mathrm{I255d/+}}$ - NT) | n |
|---|---|---|---|---|---|
| Fig. 4B | P12 - 18 | WT - NT | 16.46 ± 0.91 | $p = 0.1591$ | 5 |
| | | $R^{\mathrm{I255d/+}}$ - NT | 13.08 ± 1.40 | | 12 |
| | | $R^{\mathrm{I255d/+}}$ - CN03 | 15.70 ± 1.02 | $p = 0.0025$ | 10 |
| | | $R^{\mathrm{I255d/+}}$ - CN238 | 15.65 ± 1.24 | $p = 0.0189$ | 6 |
| Fig. 4D | P12 - 20 | WT - NT | 14.11 ± 1.35 | $p = 0.0120$ | 6 |
| | | $R^{\mathrm{I255d/+}}$ - NT | 11.19 ± 1.69 | | 7 |
| | | $R^{\mathrm{I255d/+}}$ - CN03 | 10.98 ± 1.91 | $p = 0.9996$ | 6 |
| | | $R^{\mathrm{I255d/+}}$ - CN238 | 12.82 ± 1.66 | $p = 0.1356$ | 6 |
| Fig. 4F | P12 - 24 | WT - NT | 13.35 ± 1.19 | $p = 0.0028$ | 8 |
| | | $R^{\mathrm{I255d/+}}$ - NT | 9.85 ± 4.81 | | 14 |
| | | $R^{\mathrm{I255d/+}}$ - CN03 | 7.19 ± 3.95 | $p = 0.9471$ | 7 |
| | | $R^{\mathrm{I255d/+}}$ - CN238 | 13.18 ± 2.65 | $p = 0.0656$ | 17 |
| Fig. 4H | P12 - 28 | WT - NT | 12.09 ± 1.42 | $p < 0.0001$ | 9 |
| | | $R^{\mathrm{I255d/+}}$ - NT | 3.29 ± 3.50 | | 7 |
| | | $R^{\mathrm{I255d/+}}$ - CN03 | 3.86 ± 2.53 | $p = 0.7627$ | 12 |
| | | $R^{\mathrm{I255d/+}}$ - CN238 | 9.97 ± 2.13 | $p < 0.0001$ | 13 |

**Table 7.** Effect of cGMP analogues on calpain-2 activation (calpain-2 staining) in treatments lasting 4-14 days: Quantitative data for graphs presented in Figures S1B, D, F, H.

| Figure | Cultured schema | Genotype - treatment | Mean ± SD (%) | *p* - value (compared with $R^{I255d/+}$ - NT) | n |
|---|---|---|---|---|---|
| Fig. S1B | P12 - 18 | WT - NT | 1.05 ± 0.31 | $p$ = 0.0003 | 5 |
| | | $R^{I255d/+}$ - NT | 2.93 ± 1.02 | | 9 |
| | | $R^{I255d/+}$ - CN03 | 1.92 ± 0.49 | $p$ = 0.0008 | 10 |
| | | $R^{I255d/+}$ - CN238 | 1.77 ± 0.26 | $p$ = 0.0323 | 6 |
| Fig. S1D | P12 - 20 | WT - NT | 1.33 ± 0.34 | $p$ = 0.0093 | 3 |
| | | $R^{I255d/+}$ - NT | 3.33 ± 1.44 | | 9 |
| | | $R^{I255d/+}$ - CN03 | 1.80 ± 0.71 | $p$ = 0.0077 | 8 |
| | | $R^{I255d/+}$ - CN238 | 2.18 ± 0.94 | $p$ = 0.0408 | 6 |
| Fig. S1F | P12 - 24 | WT - NT | 1.19 ± 0.34 | $p$ = 0.1385 | 9 |
| | | $R^{I255d/+}$ - NT | 1.83 ± 0.94 | | 12 |
| | | $R^{I255d/+}$ - CN03 | 1.35 ± 0.96 | $p$ = 0.3348 | 7 |
| | | $R^{I255d/+}$ - CN238 | 1.41 ± 0.58 | $p$ = 0.3904 | 12 |
| Fig. S1H | P12 - 28 | WT - NT | 0.83 ± 0.22 | $p$ = 0.0126 | 9 |
| | | $R^{I255d/+}$ - NT | 2.17 ± 1.39 | | 8 |
| | | $R^{I255d/+}$ - CN03 | 1.43 ± 0.76 | $p$ = 0.0885 | 14 |
| | | $R^{I255d/+}$ - CN238 | 1.52 ± 0.84 | $p$ = 0.1098 | 12 |

**Table 8.** Effect of cGMP analogues on calpain, HDAC, and PARP activity, PAR accumulation, after 6-day treatment: Quantitative data for graphs presented in Figures S2B, D, F, H, J.

| Figure | Genotype - treatment | Mean ± SD (%) | *p* - value (compared with $R^{I255d/+}$ - NT) | n |
|---|---|---|---|---|
| Fig. S2B calpain activity | WT - NT | 0.92 ± 0.22 | $p < 0.0001$ | 6 |
| | $R^{I255d/+}$ - NT | 2.79 ± 0.96 | | 6 |
| | $R^{I255d/+}$ - CN03 | 1.33 ± 0.17 | $p = 0.0004$ | 6 |
| | $R^{I255d/+}$ - CN238 | 0.99 ± 0.27 | $p < 0.0001$ | 7 |
| Fig. S2D HDAC activity | WT - NT | 0.97 ± 0.04 | $p = 0.1893$ | 3 |
| | $R^{I255d/+}$ - NT | 1.68 ± 0.27 | | 5 |
| | $R^{I255d/+}$ - CN03 | 1.47 ± 0.51 | $p = 0.9487$ | 7 |
| | $R^{I255d/+}$ - CN238 | 1.76 ± 0.57 | $p = 0.9771$ | 6 |
| Fig. S2F PARP activity | WT - NT | 0.93 ± 0.24 | $p = 0.0027$ | 6 |
| | $R^{I255d/+}$ - NT | 2.39 ± 1.14 | | 5 |
| | $R^{I255d/+}$ - CN03 | 1.57 ± 0.32 | $p = 0.0890$ | 6 |
| | $R^{I255d/+}$ - CN238 | 0.99 ± 0.22 | $p = 0.0037$ | 6 |
| Fig. S2H PAR accumulation | WT - NT | 0.50 ± 0.10 | $p = 0.0019$ | 3 |
| | $R^{I255d/+}$ - NT | 1.64 ± 0.50 | | 9 |
| | $R^{I255d/+}$ - CN03 | 1.39 ± 0.53 | $p = 0.6181$ | 10 |
| | $R^{I255d/+}$ - CN238 | 1.19 ± 0.32 | $p = 0.0478$ | 6 |
| Fig. S2J cleaved caspase-3 | WT - NT | 0.29 ± 0.09 | $p = 0.0039$ | 5 |
| | $R^{I255d/+}$ - NT | 1.46 ± 0.83 | | 10 |
| | $R^{I255d/+}$ - CN03 | 1.01 ± 0.63 | $p = 0.1067$ | 5 |
| | $R^{I255d/+}$ - CN238 | 1.33 ± 0.33 | $p = 0.3975$ | 5 |
